# TRAIL (CD253) Sensitizes Human Airway Epithelial Cells to Toxin-Induced Cell Death

**DOI:** 10.1101/322479

**Authors:** Yinghui Rong, Jennifer Westfall, Dylan Ehrbar, Timothy LaRocca, Nicholas J. Mantis

## Abstract

Inhalation of ricin toxin is associated with the onset of acute respiratory distress syndrome (ARDS), characterized by hemorrhage, inflammatory exudates, and tissue edema, as well as the near complete destruction of the lung epithelium. Here we report that the Calu-3 human airway epithelial cell line is relatively impervious to the effects of ricin, with little evidence of cell death even upon exposure to microgram amounts of toxin. However, the addition of exogenous soluble TNF-Related Apoptosis Inducing Ligand (TRAIL; CD253) dramatically sensitized Calu-3 cells to ricin-induced apoptosis. Calu-3 cell killing in response to ricin and TRAIL was reduced upon the addition of caspase-8 and caspase-3/7 inhibitors, but not caspase 9 inhibitors, consistent with involvement of extrinsic apoptotic pathways in cell death. We employed nCounter Technology to define the transcriptional response of Calu-3 cells to ricin, TRAIL, and the combination of ricin plus TRAIL. An array of genes associated with inflammation-and cell death were significantly up regulated upon treatment with ricin toxin, and further amplified upon addition of TRAIL. Of particular note was IL-6, whose expression in Calu-3 cells increased 300-fold upon ricin treatment and more than 750-fold upon ricin and TRAIL treatment. IL-6 secretion by Calu-3 cells was confirmed by cytometric bead array. Based on these finding, we speculate that the severe airway epithelial cell damage observed in animal models following ricin exposure is a result of a positive feedback loop driven by pro-inflammatory cytokines like TRAIL and IL-6.

## INTRODUCTION

NATO’s Biomedical Advisory Council recently concluded that ricin ranks at the top of the list of potential biothreat agents, due in large part to the toxin’s extreme potency against numerous different cell types, as well as its capacity to be disseminated via aerosol ^1^. In rodents, swine, and non-human primates (NHPs), inhalational ricin exposure evokes what is clinically equilavent to acute respiratory distress syndrome (ARDS) ^2–4^ In rodents and NHPs, the lethal dose 50 (LD_50_) of ricin by aerosol is ~4 μg/kg ^5–7^ The hallmarks of ricin-induced lung damage include early onset of alveolar macrophage apoptosis (6-12 h) followed hours later by intra-alveolar edema, accumulation of inflammatory cytokines in the BAL, neutrophilic infiltration, and fibrinous exudate^5, 7–10^. Airway epithelial cells are also a primary target of ricin intoxication and may play a role in amplifying toxin-induced pathology through secretion of pro-inflammatory cytokines and chemokines ^5, 7, 11, 12^.

Ricin itself is a potent inducer of apoptosis ^13^. The toxin is derived from castor beans (*Ricinus communis*) where it accumulates in storage vesicles as a mature, 65 kDa glycosylated protein ^14–16^. Ricin’s two subunits, RTA and RTB, are joined by a single disulfide bond. RTB is a galactose-and N-acetylgalactosamine (Gal/GalNAc)-specific lectin that promotes ricin attachment to cell surface glycoproteins and glycolipids and facilitates ricin’s retrograde transport to the endoplasmic reticulum (ER). RTA is an RNA N-glycosidase (EC 3.2.2.22) that catalyzes the hydrolysis of a conserved adenine residue within the sarcin/ricin loop (SRL) of 28S rRNA ^13, 17, 18^. In the ER, RTA is liberated from RTB and is retrotranslocated via the Sec61 complex into the cytoplasm where it inactivates ribosomes with great efficiency ^17–19^. Programmed cell death in alveolar macrophages and primary human bronchial epithelial cells occurs via caspase-3-depdendent mechanisms, although the exact upstream signals (e.g., ribsome inactivation, ribotoxic stress, MAPK and Nfκb signaling) are yet to be elucidated.

The goal of the current study was to better define the response of airway epithelial cells to the effects of ricin, especially in light of recent quantitative analysis of ribosomal depurination status that indicated that the pulmonary epithelial cells are disproportionately affected by ricin following intranasal challenge ^11^. Specifically, we reasoned that cells already compromised by ricin would be hypersensitive to secondary insults such as pro-inflammatory cytokines that are known to accumulate in the bronchoalveolar lavage (BAL) of animals following a ricin inhalation, especially the early response cytokines IL-1 and TNF-α ^2, 8, 12, 20–24^. TNF-α and related cytokines like TRAIL (tumor necrosis factor-related apoptosis-inducing ligand; CD253) are particularly suspected as having a role in driving lung epithelial cell death considering their established capacities to trigger extrinsic apoptotic cell death in different cell types experiencing intracellular stress from another insult ^25^.

## MATERIALS AND METHODS

**Chemicals and biological reagents.** Ricin toxin (Ricinus communis agglutinin II) was purchased from Vector Laboratories (Burlingame, CA). Ricin was dialyzed against PBS at 4°C in 10,000 MW cutoff Slide-A-Lyzer dialysis cassettes (Pierce, Rockford, IL), prior to use in cytotoxicity studies. Fetal calf serum was purchased from Gibco-Invitrogen (Carlsbad, CA). Cells were maintained in a humidified incubator at 37°C with 5% CO_2_. Recombinant human Tumor Necrosis Factor-α (TNF-α), recombinant human sTRAIL/Apo2L/CD253, anti-human sTRAIL-(s)-Apo2L were purchased from Peprotech (Rocky Hill, NJ). Human TNF-α neutralizing rabbit Ab was purchased from Cell Signaling Technology (Danvers, MA). Z-LEHD-FMK, Z-VAD-FMK, Z-DEVD-FMK, and Z-IETD-FMK were purchased from ApexbBio (Taiwan). Necrostatin-1 (Nec-1), GSK’872, Necrosulfonamide (NSA) were purchased from EMD Millipore (Burlington, MA). Unless noted otherwise, all other chemicals were obtained from MilliporeSigma (St.Louis, MO). Murine mAbs against ricin toxin’s A subunit (PB10, SyH7, GD12, and IB2) and B subunit (24B11, SylH3, MH3, 8A1, 8B3, LF1, and LC5) were purified by Protein A chromatography at the Dana Farber Cancer Institute (DFCI) Monoclonal Antibody Core facility (Boston, MA).

**Cell culture.** The human non-small cell lung cancer cell line (Calu-3) was obtained from American Type Culture Collection (Manassas, VA) and cultured in Eagle’s Minimum Essential Medium (EMEM) supplemented with 10% fetal bovine serum, provided by the Wadsworth Center Media Services facility. Cells were grown in a humidified incubator containing 5% CO2 and 95% air at 37°C. The cells were plated in 75 cm^2^ cell culture flasks and subcultured at 70%-90% confluence using a 0.25% trypsin solution in EDTA (Corning Life Sciences, Corning, NY). The culture medium was changed every 3 days. The cells were split 1:5 during each passage. The passages used for the following experiments were under 10.

**Cytotoxicity assay.** Calu-3 cells were trypsinized, adjusted to 5 × 10^5^ cells per ml, and seeded (100 μl/well) into 96-well plates (Corning Life Sciences, Corning, NY), and incubated for 3-4 days until confluence. Calu-3 cells were then treated with ricin, TRAIL, a mixture of ricin and TRAIL mixture, or medium alone (negative control) for 24 hr. The cells were washed to remove non-internalized toxin or TRAIL, and were then incubated for 24-72 h. Cell viability was assessed using CellTiter-GLO reagent (Promega, Madison, WI) and a Spectramax L Microplate Reader (Molecular Devices, Sunnyvale, CA). All treatments were performed in triplicate and repeated at least 3 times. 100% viability was defined as the average value obtained from wells in which cells were treated with medium only.

Based on cytotoxicity results from the ricin and TRAIL treatment above, ricin (0.25 μg/ml) and TRAIL (0.1 μg/ml) were used in all the subsequent experiments. To measure the neutralizing activity of ricin specific mAbs and neutralizing anti-TRAIL Ab, the mAbs (starting at 15 μg/ml) or anti-TRAIL Ab (starting at 1 μg/ml) in 2-fold serial dilution were mixed with ricin and TRAIL and then administered to the cells seeded in 96-well plates for 24h. After washing and then incubating for 3 days, cell viability was assessed using CellTiter-GLO reagent. Cell viability was normalized to cells treated with medium only. The protective effect of caspase inhibitors (Z-VAD-FMK, Z-LEHD-FMK, Z-DEVD-FMK, Z-IETD-FMK), RIPK1 inhibitor (Nec-1), RIPK3 inhibitor (GSK’872), and MLKL inhibitor (NSA) was also evaluated when combined with ricin and TRAIL using concentrations and incubation times as indicated in each experiment. DMSO was used as a control vehicle for all experiments.

**Caspase3/7 activity assay.** For the quantification of caspase 3/7 activities after treatment, Calu-3 cells were labeled with 500 nM Cell Event caspase-3/7 green detection reagent (Invitrogen, Carlsbad, CA) for 30 minutes at 37°C in the dark. A total of 10,000 stained cells per sample were acquired and analyzed in a FACS-Calibur flow cytometer by using CellQuest Pro software (Becton Dickinson, Franklin Lakes, NJ). Data were expressed as a percentage of total cells.

**Multiplex gene expression analysis using NanoString.** Calu-3 cells were treated with ricin, TRAIL, the combination of ricin and TRAIL, or medium alone (negative control) for 24h. RNA was extracted from treated or non-treated cells using the RNeasy plus mini kit with additional on-column DNase digestion with the RNase-Free DNase Set (Qiagen Hilden, Germany). Protocols were followed according to the manufacturer’s instructions. Extracted RNA samples were stored at −80 °C until use. Upon assay, RNA integrity was verified by agarose gel electrophoresis. RNA quality and concentration were measured using an Agilent 2100 Bioanalyzer (Life Technologies, Carlsbad, CA). RNA (100 ng) was hybridized with a predesigned nCounter human immunology panel including 594 target genes and 15 internal reference genes. The geNorm algorithm was used to select the most stable of these reference genes (GAPDH, PPIA, G6PD, EEF1G, GUSB, HPRT1, SDHA, RPL19) for normalization ^26^. The experimental procedures were carried out on the NanoString preparation station and digital analyzer according to manufacturer’s instructions. Two biological replicates were selected from each of the four groups for analysis.

**Cell supernatant cytokine quantification by cytometric bead array (CBA).** Cell supernatants were collected from treated Calu-3 cells. The BD CBA Human Inflammatory Cytokines kit (Becton Dickinson, Franklin Lakes, NJ) was used to quantitatively measure specific sets of cytokines: interleukin-8 (IL-8), interleukin-1β (IL-1β), interleukin-6 (IL-6), interleukin-10 (IL-10), TNF-α, and interleukin-12p70 (IL-12p70). Dilution series of human cytokine standards, included in the kit and prepared according to the manufacturer’s instructions, were included in each assay run to enable quantification. Assays were performed according to the manufacturer’s instructions: 50 μl of assay beads, 50 μl of the studied sample or standard and 50 μl of PE-labeled antibodies (Detection Reagent) were added consecutively to each sample tube and then incubated at room temperature in the dark for 3 h. Next, the samples were washed and centrifuged after which the pellet was resuspended in Wash Buffer and analyzed on the same day in a flow cytometer. Flow cytometry was performed using a four-laser BD FACSCalibur™ system utilizing BD CellQuest™ software for acquisition.

**Statistical Analyses**. Statistical analyses were carried out using GraphPad Prism 7 (GraphPad Software, San Diego, CA), as well as the nCounter Advanced analysis module (v 1.1.4) of the nSolver Analysis software (v3). Differential expression of the genes examined was determined by multivariate linear regression, with group membership chosen as the predictor variable and binding density as a confounder. All *p*-values derived from NanoString analysis were adjusted with Benjamini-Yekutieli correction for control of false discovery rate. Low count genes were omitted using the default settings in the nCounter Advanced analysis software for all analyses except the linear regression of gene expression values.

## RESULTS

**TRAIL sensitizes Calu-3 cells to ricin toxin-induced death.** Ricin is a promiscuous toxin capable of killing virtually all mammalian cell types. In most cases, >50% cell death occurs within 12-48 h of cells being exposed to nanogram amounts of ricin ^27^. We chose Calu-3 cells as a model to understand the response of human airway epithelial cells to ricin toxin. Calu-3 cells are a small cell lung adenocarcinoma line widely accepted as a model to study drug and nanomaterial interactions with pulmonary epithelium ^28–32^. Calu-3 cells have also been used as a model to assess the effects of other biothreat agents, namely botulinum toxin, on cells of the human airway ^33^.

To assess the sensitivity of Calu-3 cells to ricin toxin, confluent Calu-3 cells grown in microtiter plates were treated with a range of ricin toxin doses (>1-10 μg/ml) and then assessed for viability 72 h later. We found that Calu-3 cells were largely impervious to the effects of ricin to the point that we were unable to establish an IC_50_ value (**Figure S1**). Calu-3 cell monolayers grown on Transwell filters were similarly insensitive to ricin toxin (**Figure S2**).

We reasoned that pro-inflammatory cytokines like TNF-α, which is known to be released by alveolar macrophages in response to ricin, might sensitize Calu-3 cells to toxin-induced cell death ^8^. TRAIL (Apo2L; CD253) is another pro-inflammatory cytokine of interest, considering its role in accelerating lung epithelial cell death under conditions of ARDS ^34, 35^. We examined the viability of Calu-3 cells following treatment with a fixed amount of ricin (1 μg/ml) plus doses of TNF-α or TRAIL ranging from 0.01 ng/ml to 1000 ng/ml (**Figure 1A**). Calu-3 cells treated with this dose of ricin alone displayed ~80% viability at 72h time point. The addition of 10 ng/ml of TNF-α resulted in 50-60% cell death, although increasing amounts of the cytokine did not exacerbate ricin’s cytotoxic activity further indicating a level of TNF-α saturation. This is in contrast to TRAIL, which displayed a dose-dependent enhancement of ricin-induced Calu-3 cell death. TRAIL was significantly more potent than TNF-α in that virtually 100% cell killing was observed with ≥100 ng/ml TRAIL. To better define the degree of synergy between ricin and TRAIL, we performed checkerboard analysis across a range of ricin (0-1μg/ml) and TRAIL (0-1μg/ml) concentrations (**Figure 1B**). This analysis identified the minimal doses ricin (250 ng/ml) and TRAIL (100 ng/ml) required to achieve ~100% cell death within a 72 h period.

**Figure 1.**
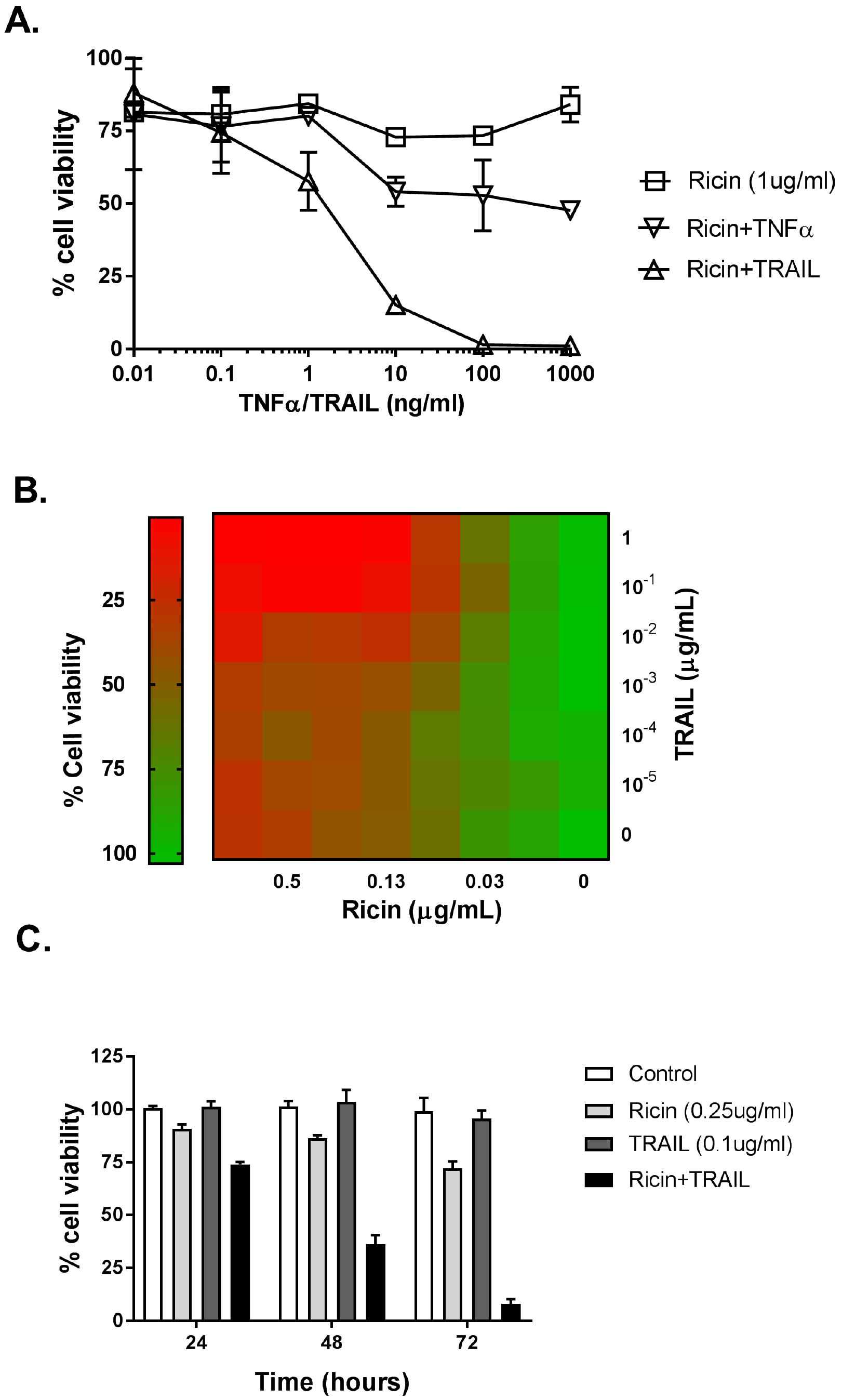
The sensitizing effect of TRAIL on ricin-induced cell death in Calu-3 cells. (A) the TNF-α or TRAIL (starting at 1μg/ml) in 10-fold serial dilution were mixed with ricin (1μg/ml) and then administrated to the cells seeded in 96-well plates for 24h. The cells were then washed and cell viability was measured 72h later, as described in material and methods. (B) In the dose experiment, cell viability was assessed at 72h after exposure to the indicated concentrations of ricin and TRAIL. (C) In time-course experiments, cell viability was assessed 24h, 48h, or 72h after cells exposed to ricin (0.25μg/ml) and TRAIL (0.1μg/ml). All treatments were performed in triplicate and repeated for 3 times. 100% viability was defined as the average value obtained from wells in which cells were treated with medium only.

A time course of Calu-3 cell viability in response to ricin (250 ng/ml), TRAIL (100 ng/ml) or the combination of ricin and TRAIL is shown in **Figure 1C**. The viability of ricin-treated Calu-3 cells declined only marginally (~25%) within a 72 h period, while viability of the TRAIL-treated cells was largely unchanged in that same time frame. In contrast, the viability of Calu-3 cells treated with the combination of ricin and TRAIL declined in a stepwise manner at 24 h, 48 h and 72 h to <25% cell viability. The observed effects of ricin and TRAIL on Calu-3 cell death was annulled by anti-human sTRAIL antibodies (**Figure 2A**) or toxin-neutralizing mAbs against RTA (**Figure 2B**) or RTB (**Figure S3**).

**Figure 2.**
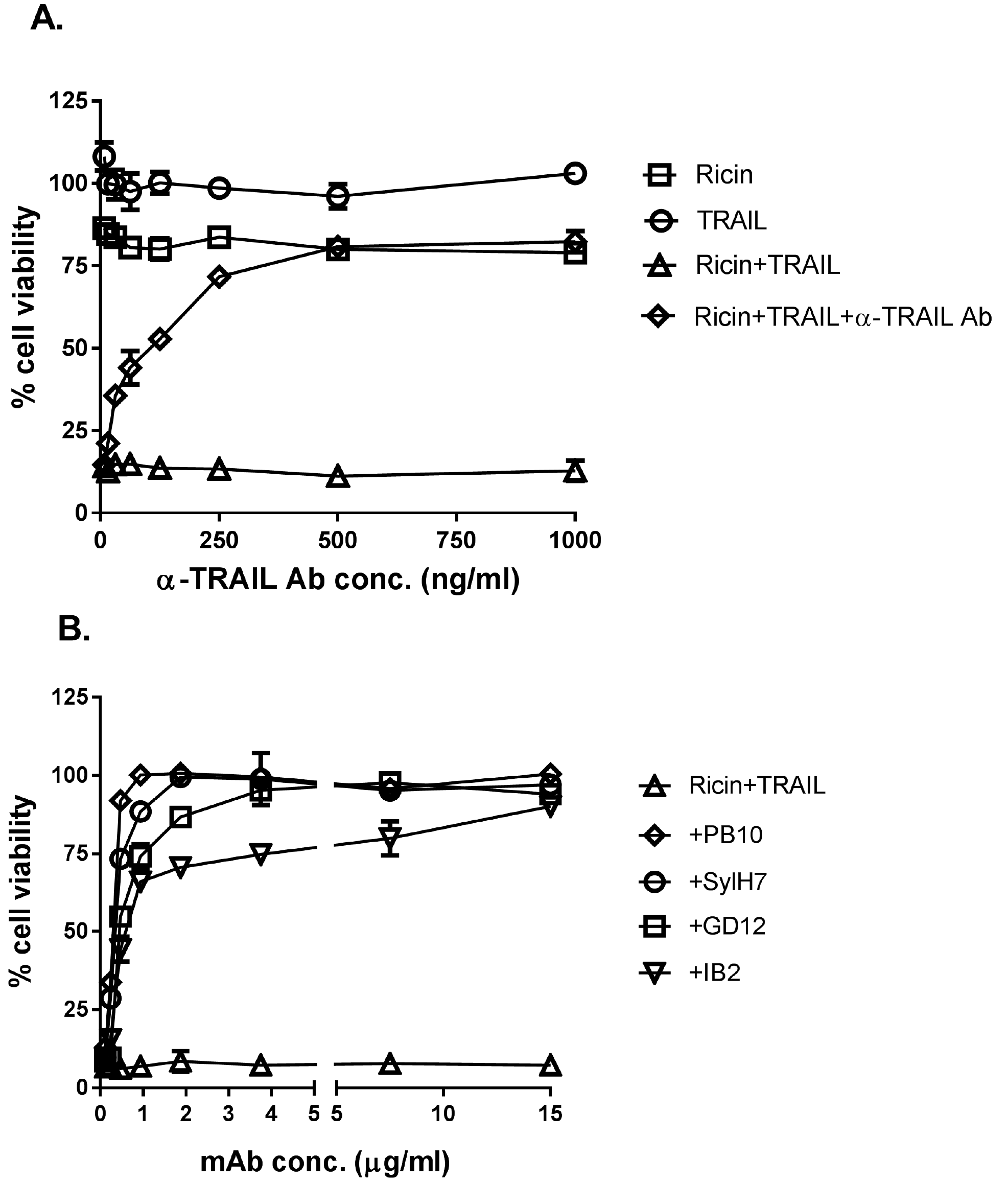
Specificity of ricin and TRAIL in inducing Calu-3 cell death. (A) anti-TRAIL ab (starting at 1 μg/ml) or (B) anti-ricin mAbs (starting at 15 μg/ml) in 2-fold serial dilution were mixed with ricin (0.25μg/ml) and TRAIL (0.1 μg/ml) and then administrated to the cells seeded in 96-well plates for 24h. The cells were then washed and cell viability was measured 72h later, as described in material and methods. The results (mean±SD) represent a single experiment done in triplicate and repeated at least three times.

It is reported that in macrophage and epithelial cells, ricin triggers the intrinsic apoptotic pathway through a process dependent on caspase-3/7 activation ^13^. We therefore examined caspase-3/7 activity in Calu-3 cells following treatment with ricin (250 ng/ml), TRAIL (100 ng/ml), or the combination of ricin and TRAIL. At the concentrations employed, neither ricin nor TRAIL alone was sufficient to induce caspase-3/7 activity in Calu-3 cells (**Figure 3A**). However, the combination of ricin and TRAIL resulted in a significant increase (~4-fold) in caspase-3/7 activity, which was inhibited by Z-DVEVD (**Figure 3A,B**). In a Calu-3 cell viability assay, ZVAD (pan-caspase inhibitor), ZIETD (caspase-8 inhibitor) and ZDEVD (caspase-3/7 inhibitor) were each able to partially suppress the cytotoxic effects of ricin and TRAIL, but only at early time points (**Figure 4**). Blocking the initiator caspase 9 with the inhibitor LEHD had no effect on Calu-3 cell viability following ricin and TRAIL (**Figure S4**), nor did treatment with the necrosis inhibitors NSA, GSK, or Nec-1 (**Figure S4**). Collectively, these results are consistent with ricin and TRAIL treatment activating apoptosis through caspase-8 and caspase 3/7-dependent pathways.

**Figure 3.**
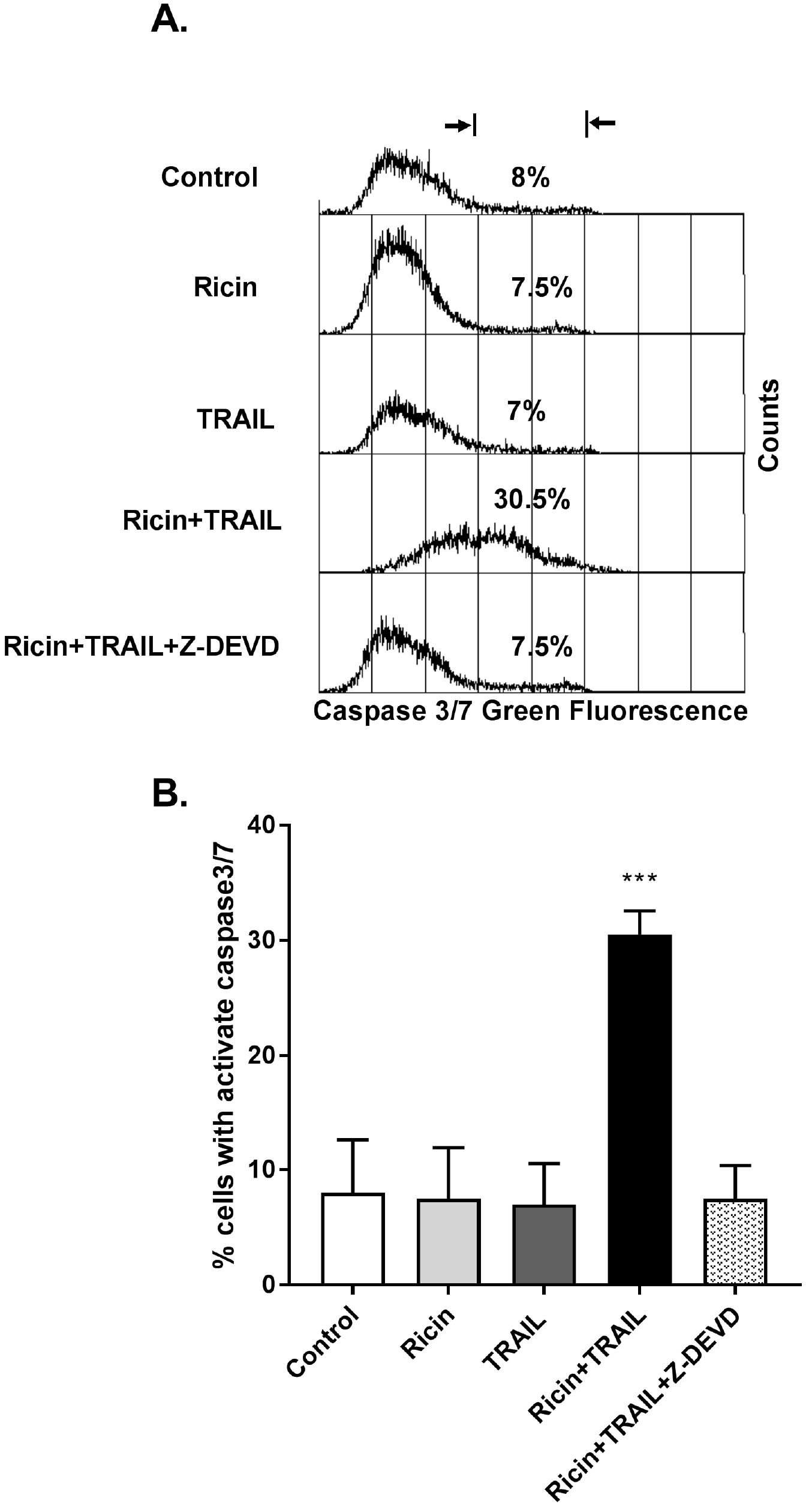
Increased caspase3/7 activity in ricin and TRAIL treated Calu-3 cells. For the quantification of caspase 3/7 activity, Calu-3 cells were treated with ricin (0.25μg/ml), TRAIL (0.1μg/ml), the mixture of ricin and TRAIL for 24h, or medium only (negative control). the caspase3/7 activity were determined by flow cytometry as described in material and methods. Caspase3/7 activity was expressed as a percentage of total cells. The results are presented as the mean±SD of three independent experiments. * p<0.01 versus control cells.

**Figure 4.**
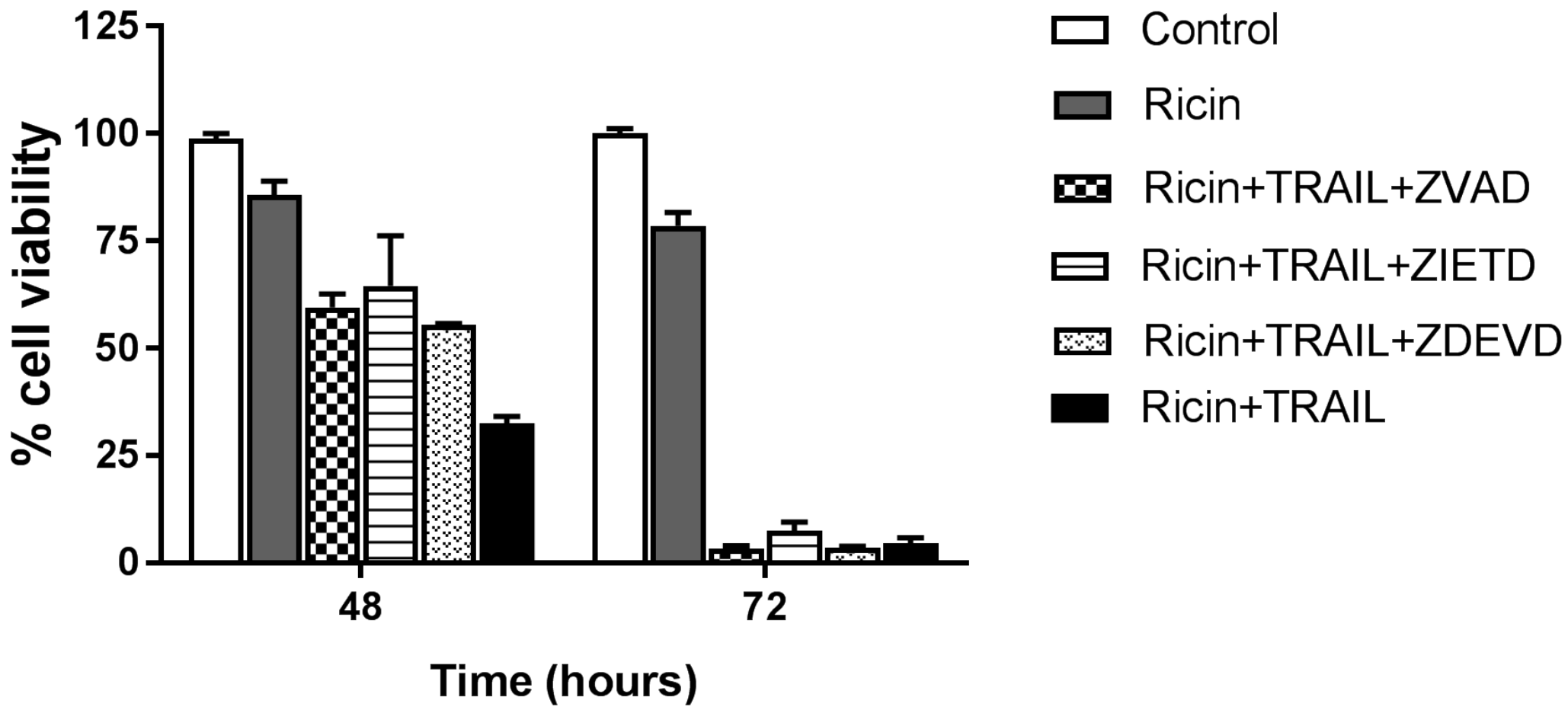
Protective effect of caspase inhibitors on cell viability in ricin and TRAIL treated Calu-3 cells. Pan-caspase inhibitor Z-VAD-FMK, Caspase 3 inhibitor Z-DEVD-FMK, or Caspase 8 inhibitor Z-IETD-FMK at 62.5nM were mixed with ricin (0.25μg/ml) and TRAIL (0.1μg/ml) and were administrated to the cells for 24h. The cells were then washed and cell viability was measured 48h or 72h later. The results (mean±SD) represent a single experiment done in triplicate and repeated at least three times.

**Transcriptional profiling of Calu-3 cells following ricin and TRAIL treatment.** To better understand the interaction between ricin and TRAIL, we subjected Calu-3 cells to transcriptional profiling using a human immunology nCounter array encompassing ~600 genes target genes. RNA was isolated from Calu-3 cells treated with ricin (250 ng/ml), TRAIL (100 ng/ml) or the combination of ricin and TRAIL for 3 h, 6 h and 18 h. At the 3 and 6 h time points, there were no significant changes in RNA levels among the target genes represented on the human immunology array when we compared ricin, TRAIL or ricin + TRAIL treatments to medium control samples (data not shown). By 18 h, the picture was markedly different. Analysis of the RNA from Calu-3 cells treated with the combination of ricin and TRAIL indicated that there were ~80 genes whose expression was elevated >2 fold over the untreated controls, which corresponds to roughly 12% of all the genes on the human immunology nCounter array (**Figure 5; Table S1; Figure S5**). Most notable was an increase in IL-6 (~750 fold), followed by other pro-inflammatory cytokines like IFN-β (~120-fold), TNF-α (~120-fold), IL-8 (88-fold) IL-1α (60-fold), CCL20 (90-fold). Virtually the same transcriptional profile was observed when Calu-3 cells were treated with just ricin, although the magnitude of the response was dampened as compared to ricin + TRAIL (**Figure 5; Table S1; Figure S5**). TRAIL treatment alone did not significantly alter Calu-3 gene expression. These results are consistent with TRAIL enhancing the pro-inflammatory and apoptotic responses of Calu-3 to ricin, rather than inducing parallel or convergent pro-inflammatory and apoptotic pathways.

**Figure 5.**
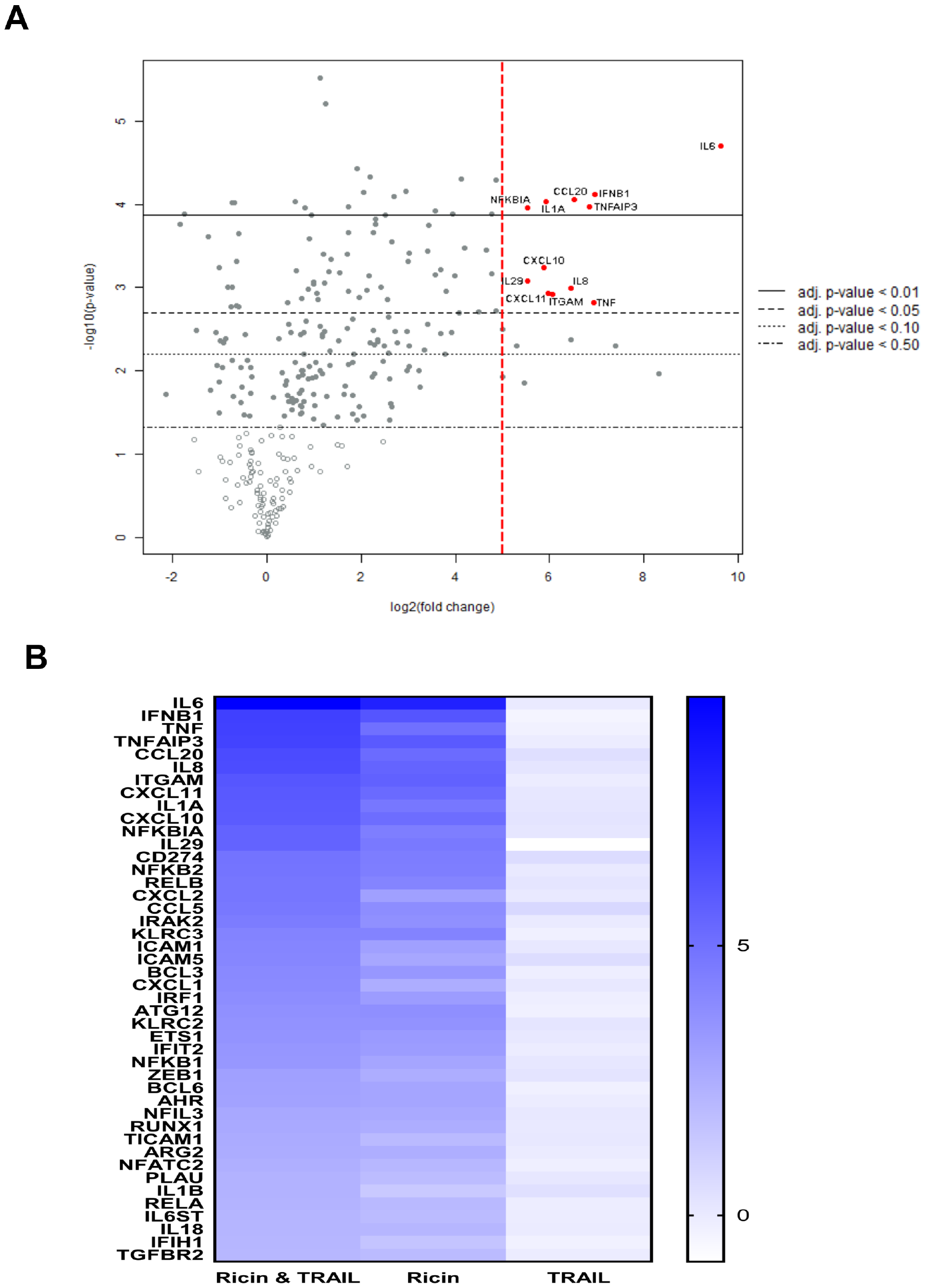
Nano-string analysis of genes differentially expressed in ricin and TRAIL treated Calu-3 cells. Calu-3 cells were treated with ricin (0.25μg/ml), TRAIL (0.1μg/ml), the mixture of ricin and TRAIL, or medium only (negative control) for 18h. RNA was extracted and subjected to nCounter analysis using the Human Immunology array panel. (**A**) Volcano plot representation of gene expression changes in ricin + TRAIL treated cells, compared with control cells. Red circles represent transcripts up-regulated >32-fold (5 log^2^). The vertical dashed red line marks the 5 log^2^ fold change threshold. **(B)** Heat map showing the relative fold change of selected genes in each treatment group compared to control. Genes were selected based on a minimum 4-fold (2 log^2^) higher expression between control and ricin + TRAIL treated cells. The color scale bar denotes maximum counts in **blue** and minimal counts in **white**. Actual fold changes (relative to control cells) are shown in **Table S1**.

To validate the transcriptional profiling studies, Calu-3 cells were treated for 24 h with ricin, TRAIL or the ricin + TRAIL combination, after which culture supernatants were assayed for the pro-inflammatory cytokines IL-6, IL-8, IL-1, IL-10, TNF-, and IL-12 levels by CBA. We found that IL-8, IL-1, IL-10, TNF-α, and IL-12 levels were unchanged, irrespective of whether Calu-3 cells were treated with ricin, TRAIL, or the ricin + TRAIL combination (**Figure 6**). IL-6 levels, in contrast, were elevated >10 fold in supernatants from Calu-3 cells treated with ricin + TRAIL, as compared to controls (**Figure 6**). Treatment of cells with ricin alone enhanced IL-6 levels, although not to a degree that was statistically significant. Thus, IL-6 expression was optimally induced by ricin + TRAIL, whereas levels of IL-8, IL-1, IL-10, TNF-α, and IL-12 were unchanged under these conditions even though corresponding mRNA levels were significantly enhanced, according to nCounter analysis.

**Figure 6.**
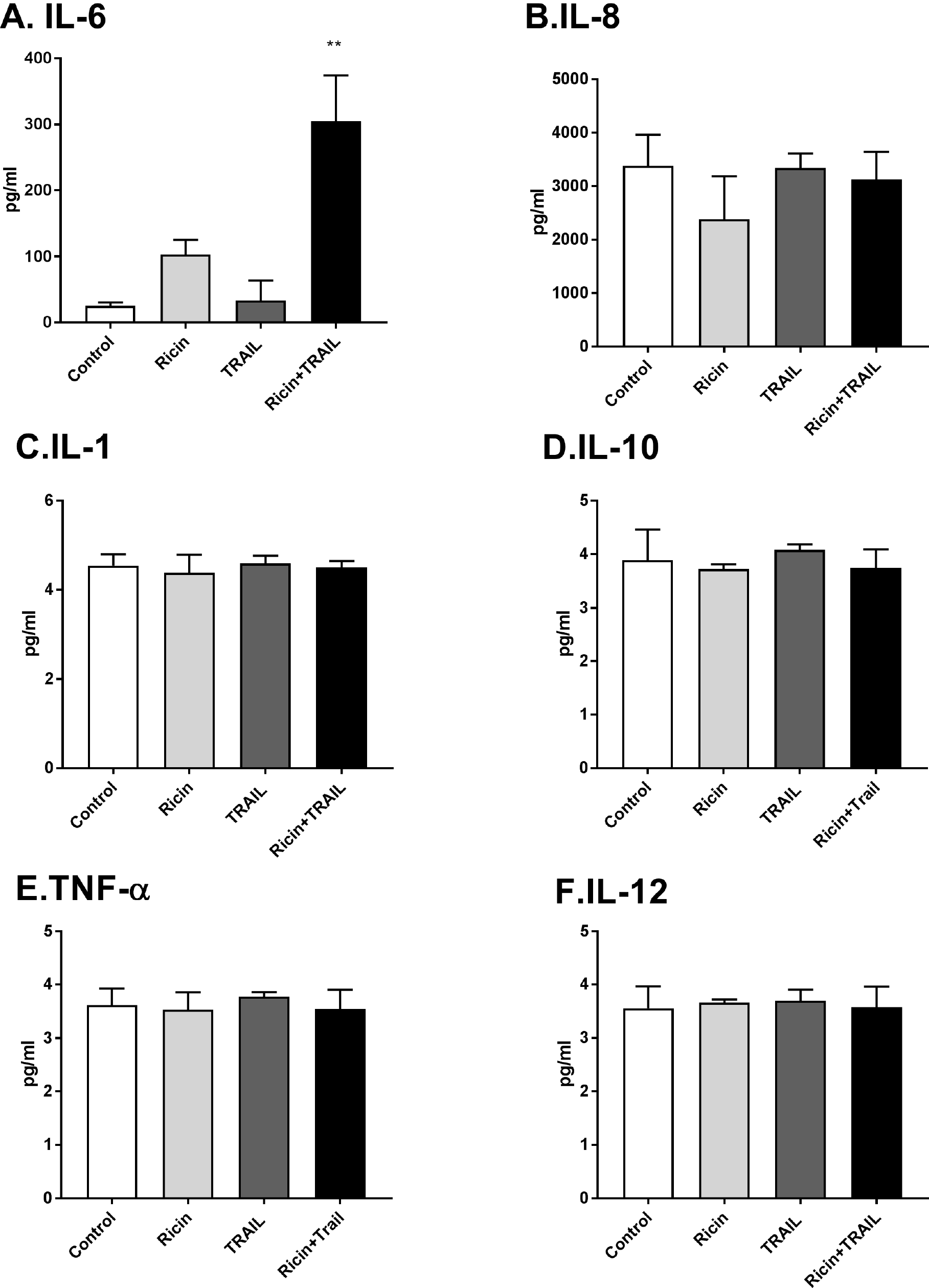
Cytokine secretion by Calu-3 cells following ricin and TRAIL treatment. Calu-3 cells were treated with ricin (0.25 μg/ml), TRAIL (0.1 μg/ml), the mixture of ricin and TRAIL, or medium only (negative control) for 24h. Cell supernatants were collected from treated cells. The levels of cytokine IL-8, IL-1β, IL-6, IL-10, TNFα, and IL-12p70 (A-F, respectively) were measured by CBA. The results are presented as the mean±SD of three independent experiments. * p<0.01 vs. untreated cells (negative control).

We postulated that the absence of TNF-α in the Calu-3 cells supernatants following ricin and TRAIL treatments might be due to autocrine signaling such that soluble cytokine is rapidly captured by TNF-α receptor, which in turn might contribute to ricin-induced cell death. To examine this possibility, Calu-3 cells were treated with ricin plus TRAIL in the presence of neutralizing anti-TNF-α antibody. We found that anti-TNF-α antibody treatment did not prevent or even reduce Calu-3 cell killing in response to ricin plus TRAIL, whereas treatment with an anti-TRAIL neutralizing antibody did rescue the cells (**Figure S6**).

## DISCUSSION

Widespread damage to the airway epithelium is a hallmark of inhalational ricin exposure, although the exact molecular events that culminate in epithelial cell destruction have not been fully elucidated ^2, 3, 5, 7, 24, 36^. In this report we utilized the well-characterized Calu-3 cell line as a prototype to better define the response of human airway epithelial cells to ricin ^28–32^. We found that Calu-3 cells, when grown to confluence on solid or permeable substrates, were largely impervious to the effects of ricin. However, co-administration of soluble TRAIL (and to a lesser degree TNF-α) rendered Calu-3 cells >1,000-fold more sensitive to toxin-induced apoptosis, as determined at 48 h and 72 h time points. At early time points, soluble TRAIL magnified the pro-inflammatory transcriptional response of Calu-3 cells to ricin, as evidenced by an across the board up regulation of genes encoding cytokines and chemokines like IL-6, TNF-α, IL-8, CCL20, and IL-1α. While the current investigation is limited to *in vitro* studies, the results are consistent with a model in which pro-inflammatory cytokines like TRAIL amplify epithelial stress-induced signal transduction pathways involved in recruitment of PMNs to the lung mucosa and, simultaneously, lowering the threshold level of ricin required to induce epithelial apoptosis.

TRAIL has previously been implicated in driving respiratory pathology and airway epithelial cell death in mice and humans in response to pathogenic agents, notably influenza virus, respiratory syncytial virus (RSV), and chlamydia ^34, 35, 37–39^. In the case of influenza virus infection, alveolar macrophages are the primary source of TRAIL ^34, 35^. Bronchial epithelial cells express TRAIL receptor(s), which ultimately modulate TRAIL-dependent apoptosis. In rodents and non-human primates, alveolar macrophages are a primary target of ricin following intranasal and inhalational challenge, so it is plausible that these cells also serve as a source of TRAIL following toxin exposure ^3, 5, 9, 40^. The availability of TRAIL neutralizing antibodies and TRAIL knock out mice will enable us to sort out the relative contribution of this cytokine to ricin-induced airway inflammation *in vivo* ^34^.

The issue of how TRAIL sensitizes Calu-3 cells to ricin-induced cell death is of particular interest, considering that TRAIL activates cell death through an extrinsic apoptotic pathway, while ricin triggers intrinsic pathways induced as a result of the ribotoxic stress response (RSR), unfolded protein response (UPR) and/or increased levels of intracellular calcium ^13, 25^. We postulate that caspase-3 serves as the central node through which ricin and TRAIL intersect. In human cells, activation of TRAIL receptor 1 (TRAIL-R1) and/or receptor 2 (TRAIL-R2) stimulates caspase 8 activation, which in turn triggers caspase-3 ^41^. Ricin-induced programmed cell death is also dependent on caspase-3 activation, although which specific upstream signaling pathway(s) (*e.g.,* RSR, UPR, ER stress) is most relevant in airway epithelial cell killing have not been completely elucidated ^2, 12^. As demonstrated in this study, Calu-3 cell death following with ricin and TRAIL treatment coincided with an increase in caspase-3/7 activity and was partially inhibited by the addition of Z-DEVD-fmk, but not impacted by LEHD, an inhibitor of caspase 9. Calu-3 cell death may also be exacerbated by a concomitant decline in endogenous inhibitors of apoptosis such as cFLIP, due to ricin’s capacity to arrest protein synthesis, as reported in the case of cells treated with Shiga toxin ^42^. In fact, the protein synthesis inhibitor cycloheximide is commonly used as a tool to sensitize cells to TRAIL-induced apoptosis ^25, 43^. While somewhat a question of semantics, we would argue that TRAIL sensitizes Calu-3 cells to ricin-induced apoptosis, rather than ricin-sensitizing Calu-3 cells to TRAIL-induced cell death. This claim is best supported by transcriptional profiling we performed which demonstrated that TRAIL amplifies across the board the effects of ricin on the treatment of Calu-3 cells. Treatment with TRAIL alone had no effect on Calu-3 gene expression, nor did TRAIL (by itself) negatively influence Calu-3 cell viability.

We identified IL-6 as being markedly up regulated in Calu-3 cells at the transcriptional and protein levels following ricin and ricin plus TRAIL treatments. This finding may have important implications for understanding the pathology associated with pulmonary ricin exposure, especially in non-human primates where elevated levels of IL-6 in bronchoalveolar lavage (BAL) fluids are associated with negative outcomes following toxin exposure (C. Roy, Y. Rong, D. Ehrbar, N. Mantis, manuscript in preparation). IL-6 also accumulates in BAL fluids and serum of mice following intranasal ricin challenge (Y. Rong and N. Mantis, manuscript in preparation) ^9, 22, 40, 44^. Whether IL-6 is more than just a biomarker of ricin intoxication remains to be determined, but there is considerable evidence to implicate this cytokine in driving local and systemic pathologies ^45, 46^. Surprisingly, Calu-3 cells did not secrete detectable levels of other “initiator” cytokines TNF-α and IL-1, or the chemokine IL-8, even though each of their respective mRNA transcripts were significantly unregulated by ricin or ricin plus TRAIL. Wong et al noted that TNF-α secretion actually declined when primary human bronchial epithelial cells were exposed to ricin, presumably due to a global arrest in protein synthesis ^12^. The continued (and possibly preferential) synthesis of IL-6, not TNF-α, in ricin intoxicated cells may have to do with differential rates of mRNA stability, especially when mitogen-activated protein kinase (MAPK) signaling pathways are activated ^47^. It is worth noting that Wong and colleagues reported that in primary human bronchial epithelial cells TNF-α and IL-1β expression in response to ricin is dependent on NfκB, but IL-6 is not ^12^.

## ACKNOWLEDGEMENTS

We gratefully acknowledge Dr. Renjie Song in the Wadsworth Center’s Biochemistry and Immunology Core facility for assisting and for the use of the flow cytometer. We thank Navjot Singh and Susan L McHale the Wadsworth Center’s Applied Genomics Technologies Core for assistance with the nCounter instrumentation. We also thank Amanda Poon (University at Albany) for helpful discussions. This work was supported by Contract No. HHSN272201400021C from the National Institutes of Allergy and Infectious Diseases, National Institutes of Health. The content is solely the responsibility of the authors and does not necessarily represent the official views of the National Institutes of Health. The funders had no role in study design, data collection and analysis, decision to publish, or preparation of the manuscript. The nCounter experiments were funded by the Director’s Office of at the Wadsworth Center in support of collaborations between the Wadsworth Center and ACPHS.

